# Distinct roles of medial prefrontal cortex subregions in the consolidation and recall of remote spatial memories

**DOI:** 10.1101/2024.03.22.586327

**Authors:** Eleonora Centofante, Mattia Santoboni, Elena L.J. Mombelli, Arianna Rinaldi, Andrea Mele

## Abstract

It is a common believe that memories with time become progressively independent of the hippocampus and are gradually stored in cortical areas. This view is mainly based on evidence demonstrating an impairing effect of prefrontal cortex (PFC) manipulations in the retrieval of remote memories paralleled by a lack of effect of hippocampal inhibition. What is more controversial is whether activity in the mPFC is required immediately after learning to initiate the consolidation process. Further question are possible functional differences among the subregions of the PFC in the formation and storage of remote memories.

To address these issues, we directly contrasted the effects of loss-of-function manipulations of the the anterior cingulate cortex (aCC) and the ventro-medial prefrontal cortex (vmPFC), that includes the infralimbic and the prelimbic cortices, before testing, and immediately after training, on the ability of CD1 mice to recall the location of the hidden platform in the Morris water maze. To this aim we injected in the vmPFC or in the aCC an AAV carrying the hM4Di receptor. Interestingly, pre-test administrations of clozapine-N-oxide (CNO) revealed that the aCC, but not the vmPFC, is necessary to recall remote spatial information. Furthermore, systemic post-training administration of CNO (3mg/kg) impaired memory recall at remote time points but not recent time points in both experimental groups. Overall, these findings revealed a functional dissociation between the two prefrontal areas, demonstrating that they are both involved in the early consolidation of remote spatial memories, but that only the aCC is engaged in their recall.

**Significant statement:** The PFC constitute an heterogeneous brain region from both a neuroanatomical and a functional point of view in which at least two main component can be distinguished: the vmPFC, with extensive connections with the limbic system and the HPC and the aCC characterized by its reciprocal connections with the sensorimotor cortex (Heidbreder and Groenewegen, 2003; Uylings et al., 2003; Le Merre et al., 2021). Contrasting the two components of the PFC, the aim of this study was twofold: investigate their relative contribution to the recall of remote spatial memory and in the consolidation of both recent and remote memories. The results demonstrated an interesting dissociation between the two components. Loss-of-function manipulation of the aCC but not the vmPFC impaired recall of remote spatial memory, however, both subdivisions contributed to its consolidation.

## Introduction

The central role of the hippocampus in episodic memory has long been established (Scoville and Milner, 1957). Nevertheless, common view is that over weeks, the role of the hippocampus progressively degrades and cortical structures become increasingly important in the stabilization of memory (Frankland and Bontempi, 2005). Two lines of evidence support this hypothesis. Firstly, mapping neuronal activity at early and late stages after training demonstrates increased (^14^C)2-deoxyglucose uptake and immediate early gene (IEG) expression in cortical areas after remote but not recent memory retrieval. Secondly, loss of functions manipulations of the prefrontal cortex (PFC) impair expression of memory at remote but not recent time points (Bontempi et al., 1999; Maviel et al., 2004; Teixeira et al., 2006; Lesburguères et al., 2011; Lopez et al., 2012; Wartman et al., 2014). Collectively, these observations have given rise to the view that the role of the neocortex is limited to the late stages of memory processing to subserve remote memory storage and recall. However, although this hypothesis has been convincingly demonstrated for aversively motivated tasks the extent to which it applies to spatial memories remains more controversial (Martin et al., 2005; Sutherland and Lehmann, 2011). Moreover, most studies investigating the engagement of the PFC in remote memory have primarily focused on recall, while much less is known about its potential involvement in the consolidation process, which could provide relevant insights into the temporal dynamics of its participation.

The relative contribution of the different components of the PFC in remote spatial memory is also a key issue. The PFC constitute a heterogeneous brain region and based on recent connectivity studies it has been suggested that at least two subdivisions of the PFC can be recognized, the dorsomedial PFC (aCC), and the ventromedial PFC (vmPFC) (Heidbreder and Groenewegen, 2003; Uylings et al., 2003; Le Merre et al., 2021). Although connectivity of the two subdivisions is not completely differentiated it has been suggested that the aCC and the vmPFC can be dichotomized based on their sensory motor and limbic hodology (Heidbreder and Groenewegen, 2003; Uylings et al., 2003; Le Merre et al., 2021). In particular the vmPFC subdivision is defined by its limbic connectivity which encompasses the hippocampal formation, the amygdala and the reciprocal connectivity with the ventral tegmental area (Parent et al., 2010; Kim et al., 2017). On the other hand the aCC is distinguished by its reciprocal connections with sensory and motor cortical regions (Groenewegen et al., 1987; Le Merre et al., 2021), the striatum and by its thalamic inputs (Vertes, 2002). This anatomical subdivision is accompanied by functional distinctions in the processing of spatial information (Sutherland et al., 1988; Kolb et al., 1994) as well as memory recall (Frankland et al., 2004; Maviel et al., 2004). Particularly relevant to our study is the finding, that IEG expression demonstrates increased neural activity in both components of the PFC when recall occurs at remote time points, while causal studies reveal task-dependent effects on remote memory recall of the two PFC components (Frankland et al., 2004; Maviel et al., 2004; Lopez et al., 2012). Thus, making it difficult to draw clear conclusions on the specific role of the vmPFC and the aCC in the formation and storage of spatial memory traces.

In this study we first asked whether the two subdivisions of the PFC could play a differential role in the recall of remote spatial memory by using chemogenetic tools, demonstrating a functional difference between the vmPFC and the aCC. In light of the difference outlined in the recall of remote spatial memory, we asked whether the same differences could also be found in the consolidation. Thus, next, we investigated the role of the two components of the PFC in the consolidation of recent and remote spatial memory. Interestingly, in this case, inhibition of both the vmPFC and the aCC had similar effects, impairing the ability to recall the platform location at remote time points.

## Materials and Methods

### Subjects

Experiments were conducted on CD1 male mice (mus musculus), 7 to 9 weeks old at the time of surgery. Animals were housed in groups of four in standard cages (26.8 x 21.5 x 14.1 cm) with food and water ad libitum under a 12h (7:00 AM – 7:00 PM) light/dark cycle. The humidity and temperature of the room were constant (21±1°C). Every experimental procedure was conducted during the light period (9:00 AM – 5:00 PM). The maximum effort was made to minimize animal suffering. Procedures were conducted under authorization N. 450-2018 form the Italian Minister of Health, according to Italian (DL 116/92) and European law regulation on the use of animals in research, and NIH guidelines on animal care.

### Stereotaxic surgery

Mice were deeply anesthetized with 3% isoflurane (Isovet; Piramal Healthcare) and secured on the stereotaxic apparatus (David Kopf Instruments) with a mouse adapter and lateral ear bars. Following craniotomies on the skull glass pipettes were lowered in the vmPFC or in the aCC according to Franklin and Paxinos (Franklin BJ and Paxinos G, 1997). Coordinates used: for the vmPFC AP: +1.7 mm, ML: ±0.3 mm, DV: -2.3 mm; aCC AP: +0.8 mm, ML: ±0.2 mm, DV: -1.8 mm, relative to bregma. Viral delivery was performed with glass micropipette connected to a syringe. Once the micropipette reached the correct placement was left in place for 1min before starting the injection. Adeno-associated viral vector (AAV2) was used to express the mCherry fluorescent protein and human influenza hemagglutinin (HA)-tagged hM4Di under the synapsin1 promoter (pAAV-hSyn-hM4D(Gi)-mCherry; Addgene #50475-AAV2). The volume injected was always 0.3 μL, for each hemisphere, over 2 minutes. After injection pipette was left in place for 5 additional minutes to allow diffusion. During surgery, the eyes of animals were maintained wet with a 0.9% saline solution and the wound was disinfected with Betadine. After surgery, the mice were placed in a recovery cage at a controlled temperature with the use of a thermal plate, before returning to the animal room for recovery.

### Behavioural apparatus and procedure

The apparatus used for the Morris Water Maze consisted of a circular pool (110 cm diameter and 40 cm high), filled up to 5 cm from the edge with black-coloured (Giotto, Italy) water (22 ± 1 °C) as previously described (Ferretti et al., 2010; Centofante et al., 2023). Black curtains surrounded the pool, and several visual cues were attached on them at the distance of round 50 cm from the pool. Four starting positions were located equidistantly around the edge of the wall, dividing the swimming pool into four equal quadrants. During the training, a circular black goal platform (10 cm of diameter), covered with wire mesh to avoid slipping, was positioned 18 cm from the wall. The apparatus was illuminated by a white diffuse light.

The procedure consisted of three different phases: a familiarization phase (on day 1), a training phase (on day 2), and one or two probe tests: 30 days or 24 hours and 30 days after the last training session respectively for the recall or the consolidation experiments. Before each phase animals were placed in isolation cages, without food or water, for 30 mins. Each session during familiarization and training consisted of three different 60 s trials, while the probe test consisted of a single 60 s trial. The familiarization phase consisted of 3 trials (intertrial interval: 20s) during which the extra maze cues were removed and the platform was made visible (1 cm over the water). During the training phase mice were submitted to 6 consecutive sessions (intersession interval: 10-15 min) of three trials (intertrial interval: 30sec). During the probe test, the platform was removed, and mice were allowed a 60s search for the platform starting from the centre of the pool. At the end of the behaviour procedures mice were perfused to verify the viral expression. All trials were recorded by a camera located over the pool and they were acquired and analysed using an automated video tracking system (Anymaze 5.0, Stoelting).

### Drug injection

Clozapine-N-oxide (CNO; HelloBio) 3 mg/Kg was dissolved in saline 0.9% and made fresh every other week. For the experiment that intended to investigate the role of vmPFC or aCC in the consolidation of remote spatial memory, CNO was administered immediately after the last training session, for experiments that intended to investigate the role of vmPFC or aCC in the retrieval of remote spatial memory it was administered 30 min before test.

### Viral detection

For mCherry fluorescent protein detection procedure mice were deeply anesthetized with chloral hydrate 5 mg/Kg and transcardially perfused with 40 ml of saline 0.9% solution followed by 40 ml of paraformaldehyde solution 4% in PBS (PFA). Brains remained in PFA 4% for 24 h before being maintained in sucrose solution 30% for others 24 h. 40 μm coronal sections were cut using a freezing microtome (Leica Microsystem, Germany) and mounted on slides cover slipped with Fluoromount with dapi (Sigma-Aldrich, Italia). Slices were mounted once every 80 μm and mCherry fluorescent protein expression was detected under a microscope (Leica DMI6000). Microphotographs were taken at 2.5X magnification of each section where mCherry fluorescent signal was seen.

Mice were included in the experimental group of interest only when mCherry fluorescent signal was inside the target location for at least 70% of the total surface. The analysis was made in the slice with the most diffuse signal, by means of area (mm^2^) of fluorescence diffusion. Moreover, we calculated the total surface of viral diffusion, to exclude animals that presented more than 30% of the total fluorescent signal inside the adjacent cortical regions. To this aim we assigned a bregma (β) value according to the mouse brain atlas (Franklin BJ and Paxinos G, 1997). Thereafter we averaged the β of maximal expression for each experiment (consolidation and retention). In the same slice, we also calculated the mediolateral (ML) and dorsoventral (DV) extension of the signal. As before, we averaged that value for all the mice in each experimental condition to obtain a mean value of viral extension. Finally, we calculated the extension of viral infection in the anteroposterior (AP) axis by acquiring the first and the last slice in which we observed the fluorescence signal (Fig. 1C-F).

**Figure 1.**
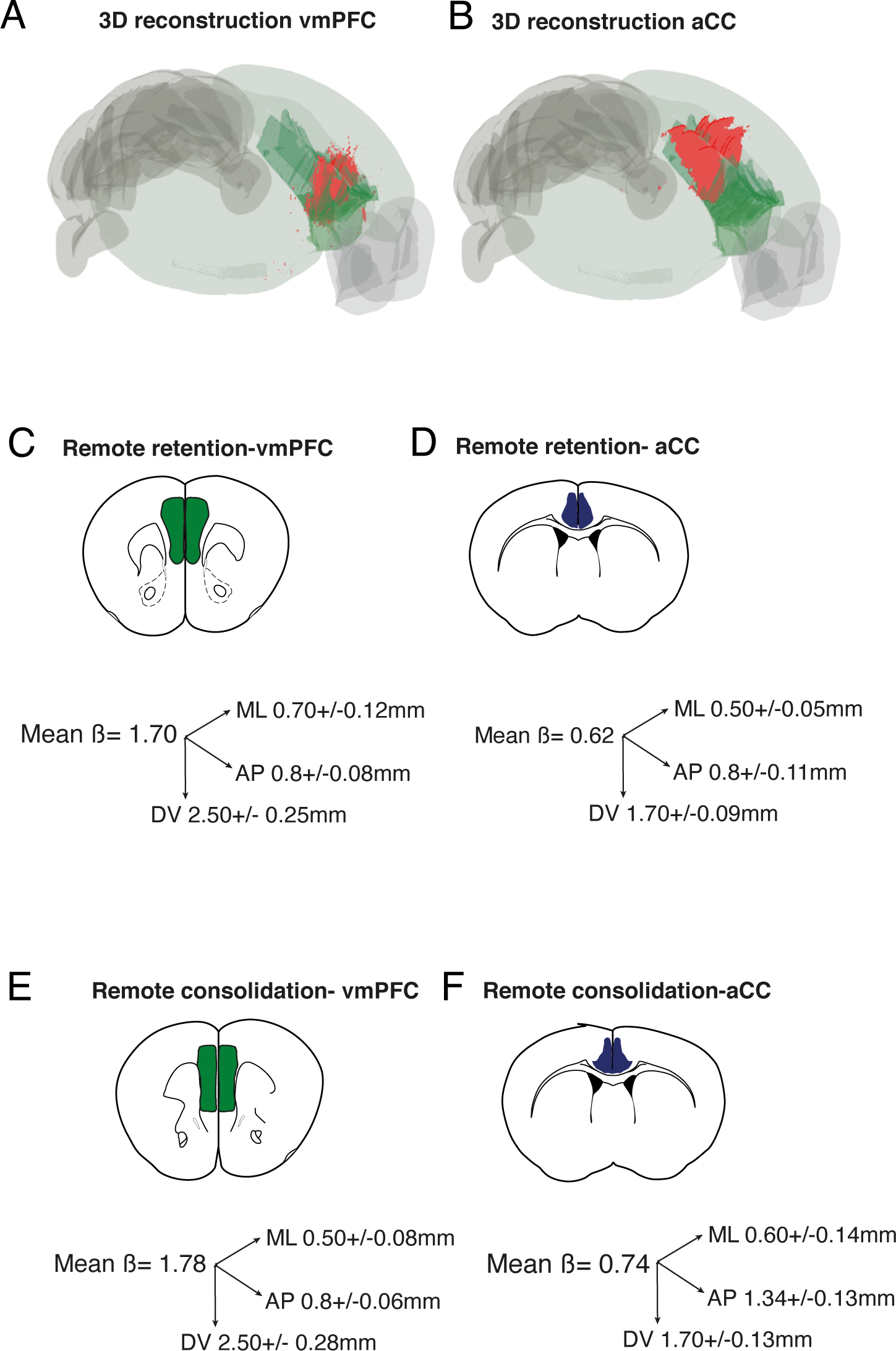
Extension of the fluorescent signal in the vmPFC and in the aCC in the different experiments. 3D reconstruction of the fluorescent in a representative mouse from the vmPFC (**A)** and from the aCC experiment (**B**). Schematic representing the fluorescent signal, and the extension of the labelling on the three axes for the different experiments. (**C)** vmPFC and **(D)** aCC labelling in the recall study; (**E)** vmPFC and **(F)** aCC labelling in the consolidation experiments. β indicates the AP coordinate (distance from bregma accordingly to (Franklin BJ and Paxinos G, 1997)) of the maximal extension of the fluorescent labeling; number on the axes indicate the average extension of the labeling on the mediolateral (ML), anteroposterior (AP), and dorsoventral axes (mean ± S.E.M.) for the animals included in the different groups.

These analyses were performed with the use of ImageJ (NIH). To create the 3D representation of the viral diffusion we used one representative animal per group (consolidation experiment and retention experiment) and applied the QUINT protocol workflow for 3D images (Yates et al., 2019) (Fig. 1A, B).

### Data collection and statistical analysis

All data are represented as mean ± standard error (SEM). MWM experiments were analysed using a two-way repeated measure ANOVA with two levels: vehicle and CNO as between groups factor and sessions or quadrants (the six sessions for the training and quadrants for the probe test) as repeated measures. When the interaction between factors was not significant, a one-way repeated measures ANOVA with the six sessions or the four quadrants as repeated measures was used on each group independently. Tukey Honest Significant Difference (HSD) was used as a post-hoc comparison.

## Results

### Assessment of viral infection in the vmPFC and in the aCC

The neuroanatomical classification of the vmPFC and the aCC was delineated accordingly to the Franklin and Paxinos (Franklin BJ and Paxinos G, 1997) mouse brain atlas and the Allen Brain Atlas (Wang et al., 2020). The vmPFC included the infralimbic (IL), the prelimbic (PL), and anterior component of the cingulate cortex (Cg1), the aCC included the posterior component of the Cg1 and the Cg2. As represented in the 3D reconstruction of the fluorescent signal in two representative mice from the vmPFC and aCC groups, the infected area was restricted to two distinct and non-overlapping regions (Fig. 1A, B). As shown in Fig. 1C-E extension of the fluorescent labelling in the mice included in the two vmPFC experiments was highly superimposed (Fig. 1C-E), similar fluorescent labelling was also observed when comparing spreading in the two aCC groups (Fig. 1C-F). Finally, it should be emphasised that quantification of anteroposterior viral diffusion in the vmPFC and in the aCC demonstrate no overlap between the infected areas. Supplementary figures 1 and 2 show the schematic representation of the area with the most diffuse fluorescent signal for all the mice included in the experiments.

### Functional dissociation of aCC and vmPFC in remote memory recall

To determine the role of these two different cortical modules in the recall of remote spatial memories we performed a pre-test chemogenetic inhibition of the vmPFC and the aCC 30 days after massed training in the Morris water maze (MWM) in two different groups of mice (Fig. 2A). As represented in Fig. 2B and C during training the latency to reach the platform decreased across the six sessions in a similar way in the two experiments for both experimental groups (vmPFC: two-way repeated measures ANOVA, session F(5, 100)= 13.10, p< 0.0001; treatment F(1, 20) = 0.2140, p = 0.648; session x treatment F(5, 100) = 1.772, p = 0.125; aCC: two-way repeated measures ANOVA session F(5, 85) = 12.65, p < 0.0001; treatment F(1, 17) = 0.3698, p = 0.5512; session x treatment F(5, 85) = 0.5212, p = 0.7596). The one-way repeated measures ANOVA revealed a significant effect of the session for both vehicle and CNO groups in vmPFC and aCC (vmPFC vehicle F(5, 55) = 6.732, p< 0.0001; vmPFC CNO F(5, 45) = 7.724, p < 0.0001; aCC: vehicle F(8, 40) = 3.128, p = 0.0018; aCC CNO F(5, 45)= 8.931, p < 0.0001) demonstrating that the animals were able to learn the task and there were no differences between groups before treatment (Fig. 2B, C).

**Figure 2.**
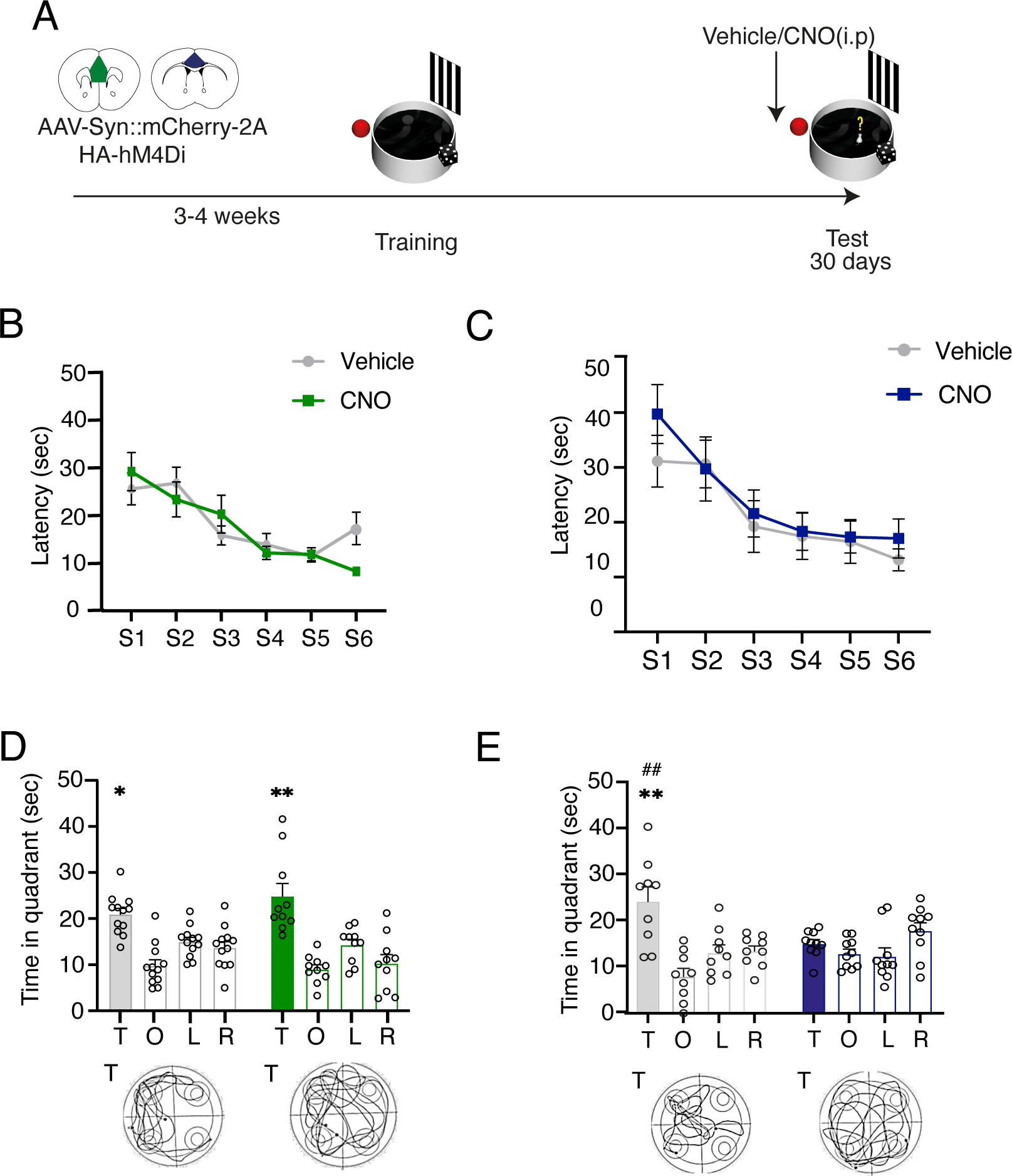
aCC but not vmPFC is required for the recall of remote spatial memories 30 days after training. Schematic of the experimental design **(A)**. Latency (± SEM) to reach the platform during training before vehicle (N= 12) or CNO (N= 10) administrations in the vmPFC infected groups **(B)**. Latency (± SEM) to reach the platform during training before vehicle (N= 9) or CNO (N= 10) administrations in the aCC infected groups **(C)**. The ability to recall the correct quadrant was not affected by pre-test systemic CNO (3mg/Kg i.p.) administrations in mice infected in the vmPFC as compared to saline **(D).** Pre-test CNO administrations impaired ability to recall the platform location as compared to saline in mice infected in the aCC **(E).** Histograms represent the mean of distance traveled in each quadrant: T= Target; O= Opposite; R= Right; L= Left. Bottom panels: representative track plots from vehicle and CNO animals in the two experiments. *p≤ 0.05; ** p≤ 0.01 vs target quadrant (within group); ## p≤ 0.01 target vs target quadrant (between groups) (Tukey, HSD).

Interestingly administrations of CNO prior to the probe test performed 30 days after training revealed a dissociation in the role of the vmPFC and the aCC in the ability to retrieve information relative to the platform location (Fig. 2D, E). As shown in Fig. 2D, mice expressing hM4Di in the vmPFC when administered before probe test with CNO i.p. where able to correctly locate the platform similarly to saline injected controls. The two-way ANOVA analysis did not highlight any differences in the time spent in the target quadrant for vmPFC groups (quadrant F(3, 60) = 23.20, p < 0.0001; treatment F(1, 20) = 0.3237, p = 0.5757; quadrant x treatment F(3, 60) = 1.556, p = 0.2094). Further analysis confirmed that both vehicle and CNO groups were able to correctly remember the platform location (one-way repeated measures ANOVA: vehicle F(3, 33) = 11.21, p < 0.0001; CNO: F(3,27) = 11.86, p < 0.0001) (Fig. 2D). These findings were confirmed by the analysis of the distance travelled (Supplementary Fig. 3A) as well as by the analysis of the annulus frequency, that is often considered directly linked to the acquisition of the correct spatial location of the platform (Ferretti et al., 2007; Lasalle et al., 2000) (Supplementary Fig. 3A).

On the contrary, pre-test CNO-injected mice in the aCC group showed an impaired ability to recall the platform location compared to control mice (two-way ANOVA repeated measures, quadrant preference F(3, 51)= 8.0, p= 0.0002; treatment F(1, 17) = 1.6, p= 0.216; quadrant x treatment F(3, 51) = 5.251, p= 0.0034) (Fig. 2E). The analysis of distance travelled and annulus frequency in the different quadrants confirmed impairment of the CNO-injected mice compared to vehicle injected controls (Supplementary Fig. 3B). The analysis on the mean speed during the probe test did not reveal any difference between the vehicle and CNO groups in the two experiments (Supplementary Fig. 3B).

Overall these results are in line with previous findings (Maviel et al., 2004; Teixeira et al., 2006) demonstrating a role of the PFC in the recall of remote spatial memory, but they also provide new insight in the functional role of the two cortical modules, demonstrating a functional dissociation between vmPFC and aCC, and establishing that only the aCC is involved in the retrieval of remote spatial memories.

### Both the vmPFC and the aCC are required for the consolidation of remote but not recent spatial memories

The establishing of stable remote memories traces requires a consolidation process that starts immediately after encoding (McGaugh, 2000). However, experimental evidence investigating the role of the PFC in the consolidation of the memory trace, based mainly on fear memory tests, is contradictory and leaves doubts on when this brain region is engaged and whether its role is limited to the consolidation of remote memories or it is also functional to the maintenance of memories at shorter intervals (Izquierdo et al., 2007; Restivo et al., 2009; Leon et al., 2010; Lesburguères et al., 2011; Bero et al., 2014; Torres-García et al., 2017; Vetere et al., 2017; Pereira et al., 2019). Therefore, next we asked whether immediate post-training loss of function manipulations of the PFC could impair consolidation of spatial memory at early and late time points by using chemogenetics. Moreover, in light of the functional dissociation between the vmPFC and the aCC found in the recall of spatial information we contrasted the two brain regions to see whether a similar dissociation could be observed also on the consolidation.

In the first experiment a group of mice, virally injected in vmPFC, was trained in the MWM and immediately post-training administered i.p. with either vehicle or CNO (Fig. 3A). No differences were found in the training of the two groups (two-way repeated measures ANOVA: sessions F(5, 70) = 14.76, p< 0.0001; treatment F(1, 14) = 0.1355, p = 0.7183; session x treatment F(5, 70) = 0.2505, p = 0.9382) that progressively reduced the time to find the platform during successive sessions (one-way repeated measures ANOVA: vehicle F(5, 35) = 6.601, p = 0.0002; CNO F(5, 35) = 9.041, p < 0.0001) confirming the ability of the mice to properly acquire the task before treatment (Fig. 3B).

**Figure 3.**
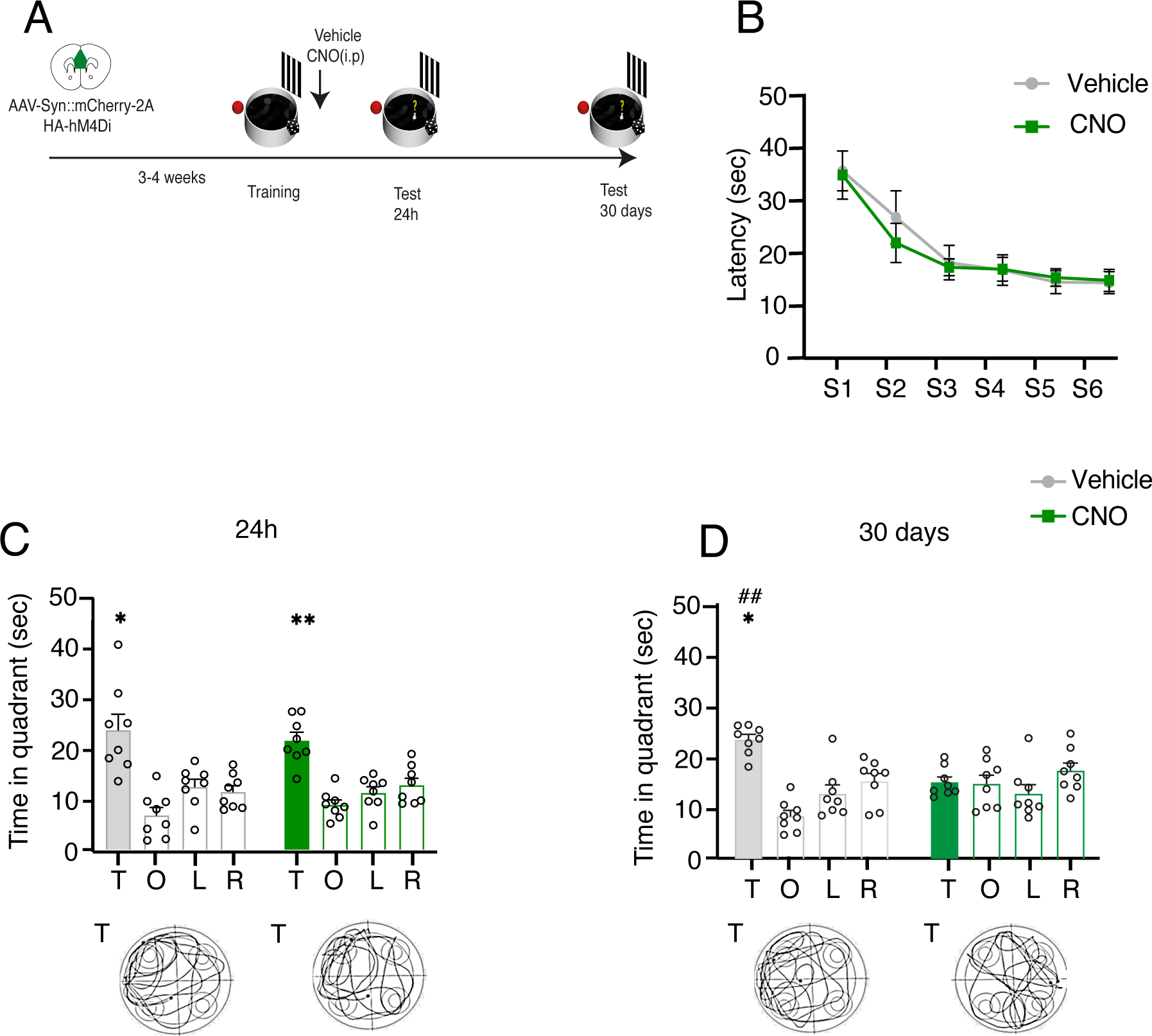
vmPFC activation is required to consolidate remote but not recent spatial memories. **(A)** Schematic of the experimental design. **(B)** Latency (± SEM) to reach the platform during training before vehicle (N= 8) or CNO (N= 8) post-training administrations in the vmPFC infected groups. Post-training systemic administrations of CNO did not affect mice ability to recall the correct platform location 24h after training compared to vehicle controls **(C)**. Mice administered post-training with CNO showed impaired performance on the probe test 30 day after training **(D)**. Histograms represent the mean of distance traveled in each quadrant: T= Target; O= Opposite; R= Right; L= Left. Bottom panels: representative track plots from vehicle and CNO animals in the two experiments. *p≤ 0.05; ** p≤ 0.01 vs target quadrant (within group); ## p≤ 0.01 target vs target quadrant (between groups) (Tukey, HSD).

When we analysed the time spent by the mice in the different quadrants during the probe test 24 hours after training, we found that both vehicle and CNO-injected mice spent significantly more time in the target quadrant (two-way repeated measures ANOVA of quadrant F(3, 42) = 21.61, p< 0.0001; treatment F(1, 14) = 1.224, p = 0.2873; quadrant x treatment F(3, 42) = 0.5290, p = 0.6648) regardless from the treatment (one-way repeated measures ANOVA vehicle F(3, 21) = 9.878, p = 0.0003; CNO F(3, 21) = 13.85, p < 0.0001) (Fig. 3C). On the other hand, CNO-injected mice showed a clear impairment in remembering the platform location 30 days after training compared to controls (two-way repeated measures ANOVA quadrant F(3, 42) = 8.788, p = 0.0001; treatment F(1, 14) = 1.690, p = 0.2146; quadrant x treatment F(3, 42) = 6.917, p = 0.0007) (Fig. 3D). The results were confirmed by the additional analyses on the distance travelled and the annulus entries either 24h and 30 days after training (Supplementary Fig. 4A, B). Once again, the results on the mean speed during the probe test confirmed that there were no differences between vehicle and CNO treated groups on general parameters of behaviour (Supplementary Fig. 4A, B).

To determine the role of the aCC in the consolidation of recent and remote spatial memory, in a second experiment we trained mice virally injected in aCC, and performed immediate post-training administrations of either vehicle or CNO to tested them 24 hours and 30 days after training (Fig. 4A). As for the vmPFC experiment we did not find any difference in the learning curve of the two groups before saline or CNO administrations (two-way repeated measures ANOVA of session F(5, 110) = 26.39, p < 0.0001; treatment F(1, 22) = 0.1514, p = 0.7009; session x treatment F(1, 22) = 0.1514, p = 0.7009). Further analysis confirmed that both groups were able to acquire information relative to the platform location, progressively reducing the time to reach the platform over successive sessions (one-way repeated measures ANOVA: vehicle F(5, 60) = 18.64, p < 0.0001; CNO F(5, 50)= 9.320, p < 0.0001) (Fig. 4B). When tested for recent memory 24 hours after training, both vehicle and CNO were able to remember the platform location (two-way repeated measures ANOVA quadrants F(3, 66) = 41.70, p < 0.0001; treatment F(1, 22) = 2.592, p = 0.1217; quadrant x treatment F(3, 66) = 0.4726, p = 0.7024), spending significantly more time in the target quadrant (one-way repeated measures ANOVA vehicle F(3,36)= 16.43, p< 0.0001; CNO F(3, 30) = 27.38, p < 0.0001) (Fig. 4C). On the contrary, when testing was performed 30 days after training, only mice administered post-training with vehicle were able to correctly recall the platform location (two-way repeated measures ANOVA of quadrant F(3, 66) = 6.549, p = 0.0006; treatment F(1, 22) = 0.5776, p = 0.4553; quadrant x treatment F(3, 66) = 2.010, p = 0.1210; one-way repeated measures ANOVA of controls F(3,36) = 7.099, p = 0.0007; CNO F(3, 30) = 1.531, p = 0.2267) (Fig. 4D). Analysis on the distance travelled and the annulus frequency highlighted the same difference between controls and CNO administered animals (Supplementary Fig. 4C, D), providing more strength to our results. Analysis of mean speed did not reveal any difference between the two. groups (Supplementary Fig. 4).

**Figure 4.**
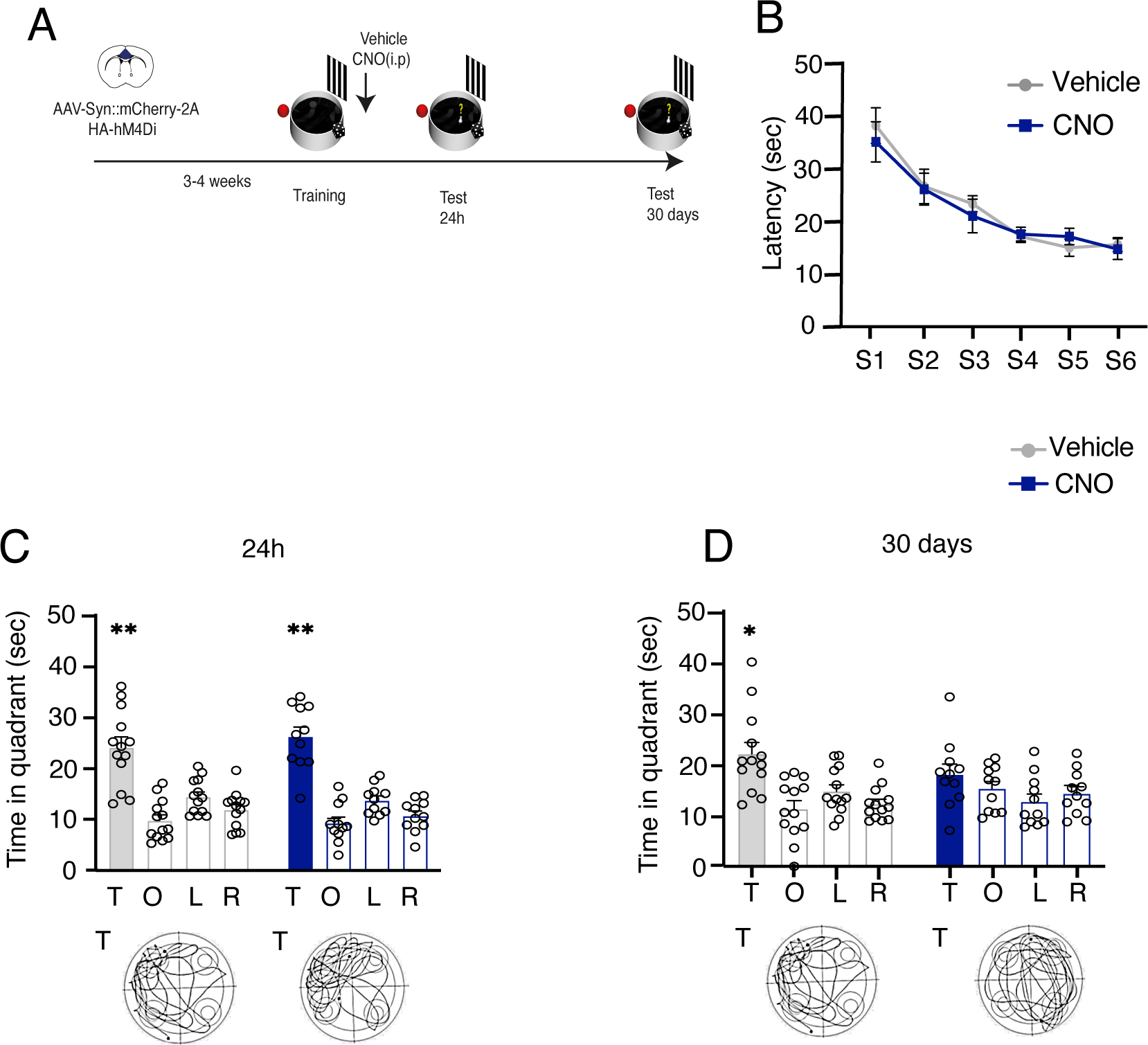
aCC activation is required to consolidate remote but not recent spatial memories. **(A)** Schematic of the experimental design. **(B)** Latency (± SEM) to reach the platform during training before vehicle (N= 8) or CNO (N= 8) post-training administrations in the aCC infected groups. Post-training systemic administrations of CNO did not affect mice ability to recall the correct platform location 24h after training **(C)**. Post-training administration of CNO impaired mice ability to recall the platform location 30 days after training. Histograms represent the mean of distance traveled in each quadrant: T= Target; O= Opposite; R= Right; L= Left. Bottom panels: representative track plots from vehicle and CNO animals in the two experiments. *p:≤ 0.05; ** p:≤ 0.01 vs target quadrant (within group) (Tukey, HSD).

These findings demonstrate that the PFC is involved in the early phases of the consolidation of remote but not recent spatial memories, moreover they revealed that the two cortical modules play a similar role in this process.

## Discussion

Contrasting the two components of the PFC, the aim of this study was twofold: investigate their relative contribution to the recall of remote spatial memory and in the consolidation of both recent and remote memories. The results demonstrated an interesting dissociation between the two components. Loss-of-function manipulation of the aCC but not the vmPFC impaired recall of remote spatial memory, however, both subdivisions contributed to its remote consolidation.

Current hypotheses on memory consolidation suggest that hours after learning molecular changes in the hippocampus lead to synaptic strengthening within this region (Kandel et al., 2014; Dudai et al., 2015). Over weeks the role of the hippocampus degrades and cortical structures become more important (Frankland and Bontempi, 2005). Our results confirm this view, demonstrating that chemogenetic inhibition of the aCC before testing impairs the ability to recall the correct platform location 30 days after the last training session. Interestingly this effect was specific to the aCC, in fact, when the same manipulation was performed in the vmPFC the animals were perfectly able to locate the platform. It should be mentioned that in both cases we performed systemic administrations of CNO, but the lack of effect in the vmPFC group exclude the possibility that the effects observed in the aCC group depend on CNO alone.

IEG expression has been found to increase after remote memory recall in both the vmPFC and in the aCC for different kind of information (Frankland et al., 2004; Maviel et al., 2004; Lopez et al., 2012). However, causal interrogation of their role in the recall of remote memory in different tasks suggest a functional distinction between the two components. For example, inactivation of the aCC impairs remote recall in both fear conditioning (Frankland et al., 2004), and in the radial arm maze (Maviel et al., 2004), on the contrary the vmPFC seems to be involved only in the recall in the radial arm maze but not contextual fear conditioning (Frankland et al., 2004; Maviel et al., 2004). Although the finding reported in the present study are in line with previous reports demonstrating the impairing effect of loss-of-function manipulation of the aCC in the recall of remote memories in the MWM (Teixeira et al., 2006; Wartman et al., 2014) to our knowledge this is the first direct comparison of the effects of loss-of-function manipulation of the two PFC component in the MWM, demonstrating a dissociation between the two PFC subdivisions similar to what has been found for contextual fear conditioning (Frankland et al., 2004). Based on the present findings only it is difficult to explain these differences. Nevertheless, it is worth noting that differences in c-fos expression the aCC and the vmPFC have been observed at remote time points in the MWM when comparing task conditions affecting memory strength (Lopez et al., 2012). Moreover, loss-of-function manipulations of the vmPFC have been shown to improve performance in the MWM at remote time points only when there is the need to solve conflicting information between new and stored information on platform location (Richards et al., 2014). These observations suggest that the vmPFC might come into play only when remotely acquired information progressively fading away need to compete for performance with strong newly acquired memories (Eichenbaum, 2017). Although very speculative this view could explain the discrepancy reported in the literature as well as our finding where there is no competition between new and remote memories in the probe trial.

In the second experiment we contrasted the two components of the PFC in the consolidation spatial memories at recent and remote time points. Surprisingly in contrast to the dissociation observed in the recall of remote memories we found that the two PFC compartments behave in a similar way. In fact, immediate post-training manipulations in either the vmPFC and the aCC impaired recall at late time points leaving unaffected the ability to correctly locate the platform 24h after training. These finding raises several interesting issues. First one is that they provide one of the first evidence that neuronal activity in the PFC immediately after training is required for the consolidation of remote but not recent spatial memory as assessed in the MWM. Studies investigating the role of the PFC in the consolidation of memory is primarily based on studies in aversively motivated tasks (Izquierdo et al., 2007; Bero et al., 2014; Torres-García et al., 2017; Vetere et al., 2017; Pereira et al., 2019). This is mainly due to the difficulty of separating the encoding and the consolidation phase in spatial learning tasks requiring training over multiple days, where consolidation occurs repeatedly at the end of each training period. In the present study we used a massed version of the MWM that allows manipulation after a single training, lasting approximately 1h, at the early stages of the consolidation phase. It should be mentioned that a similar experimental strategy has been previously adopted by Leon and colleagues (2010) reporting on the one hand learning induced increase in ERK1/2 phosphorylation in the HPC but not in the PFC immediately after training, on the other hand an impairing effect of post-training PFC administrations of the mitogen-activated protein kinase (MAPK)/ERK inhibitor (U0126) on the ability to correctly locate the platform 24 hours after training (Leon et al., 2010). Although there is an apparent discrepancy between the two observations the impairing effects induced by post-training administrations of U0126 in the PFC seem in contrast with the lack of effect we report in the present study. It is worth noting that contrasting results on the role of PFC in the consolidation of recent memory have been reported also in other learning tasks such as for example one-trial inhibitory avoidance (Izquierdo et al., 2007; Torres-García et al., 2017), fear conditioning (Bero et al., 2014; Vetere et al., 2017; Pereira et al., 2019) and odor discrimination tasks (Tronel and Sara, 2003; Lesburguères et al., 2011), depending on the PFC module manipulated, or the time window chosen for the manipulation. Further studies will be needed to better understand the contribution of PFC and its subdivisions in the consolidation of recent memories.

Another intriguing issue is the similar effect induced by loss-of-function manipulation of the two PFC components in the consolidation of remote spatial memory, despite their functional differences in remote recall. Previous studies have generally excluded early engagement of different PFC components immediately after training, based on the lack of change in neuronal activity observed through IEG expression after probe tests at early and late time points (Frankland et al., 2004; Maviel et al., 2004). Our findings support the notion that activity in both subdivisions of the PFC is required early after learning to promote the formation of stable memory traces that can be recalled at late time points. Although direct comparison between whole brain region loss-of-function manipulation such as the one we used in the present study and the engram cells approach is difficult, intriguingly utilizing activity-dependent labeling to track the activation pattern of engram cells in the PFC after training and their causal role in fear conditioning have led to similar conclusions (Kitamura et al., 2017; DeNardo et al., 2019).

What was particularly surprising to us was the dissociation observed in the vmPFC, which we found to impair the consolidation of remote spatial memories but not their recall. Therefore, interestingly, the vmPFC appears to be involved solely in the consolidation of remote spatial memory traces, differently to both the HPC (Riedel et al., 1999) and the aCC, engaged in the consolidation and recall at recent or remote time points.

The evidence presented in this study establish that neuronal activity in both the PFC subregions is necessary immediately after training in the Morris water maze to consolidate spatial information needed for correctly locating the platform at remote but not recent time points. Moreover, we demonstrated a role of the aCC but not the vmPFC in the recall of the platform location at remote time points. This suggests that after training the activity of the vmPFC gradually declines and the aCC takes over in retaining the information playing an active role in their remote recall. It could be suggested that spatial memory immediately after encoding engages a wide network of brain regions that include the HPC (Riedel et al., 1999), the vmPFC and the aCC and possibly other brain regions to initiate the consolidation process. Once the memory trace is well consolidated the information are stored over time in the aCC, then the vmPFC comes into play and participate to the recall only if there is a competition for the performance between old and new memories (Richards et al., 2014; Eichenbaum, 2017). It has been suggested that the HPC to PFC projections are important in controlling formation of stable memory traces (Barker et al., 2017). Although connectivity of the two PFC subdivisions is not completely segregated, the vmPFC is better characterized by limbic afferents projections and dopaminergic modulation (Heidbreder and Groenewegen, 2003; Uylings et al., 2003; Le Merre et al., 2021). It would therefore be intriguing to explore whether vmPFC might play a role in the selection of relevant information to be stored in the aCC gating HPC input to the PFC. Such investigations could shed further light on the complex dynamics underlying memory consolidation and recall processes.

## Acknowledgements

The authors would like thank Laura Costantini and Sara Pezza, for their help with the experiments. This study was supported by a PRIN2022 grant from MUR (to A.M.) and by grants from Sapienza University of Rome (to A.M.; A.R.).

## Notes

Conflict of interest: The authors declare no conflict of interest.

### Competing Interest Statement

The authors have declared no competing interest.

