## SupplementaryFigures for "Distinct roles of medial prefrontal cortex subregions in the consolidation and recall of remote spatial memories"

A

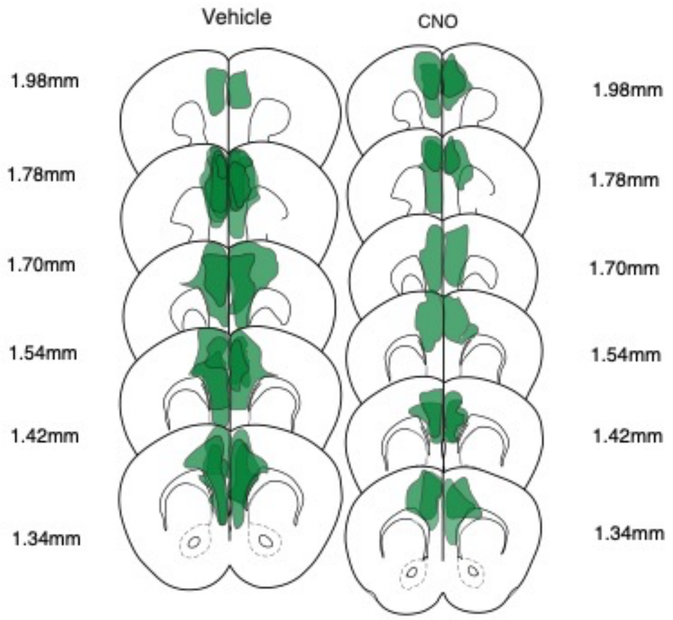
Vehicle
CNO
1.98mm
1.78mm
1.70mm
1.54mm
1.42mm
1.34mm
1.98mm
1.78mm
1.70mm
1.54mm
1.42mm
1.34mm

B

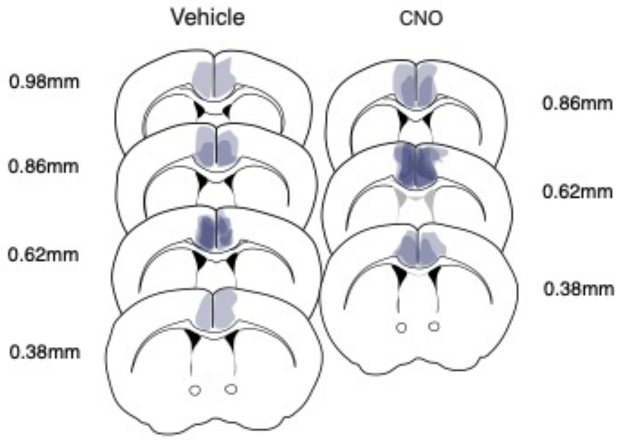
Vehicle
CNO
0.98mm
0.86mm
0.62mm
0.38mm
0.86mm
0.62mm
0.38mm

**Supplementary Figure 1. AAV-Syn::mCherry-2A-HA-hM4Di virus diffusion for the remote retention experiments.** Schematic representation of mCherry protein expression in vmPFC **(A),** and in the aCC **(B)**. mCherry maximum extension is represented in green for each single mouse for the vmPFC group and in blue for each single mouse of aCC group. Coordinates are expressed as mm anterior to bregma.

A

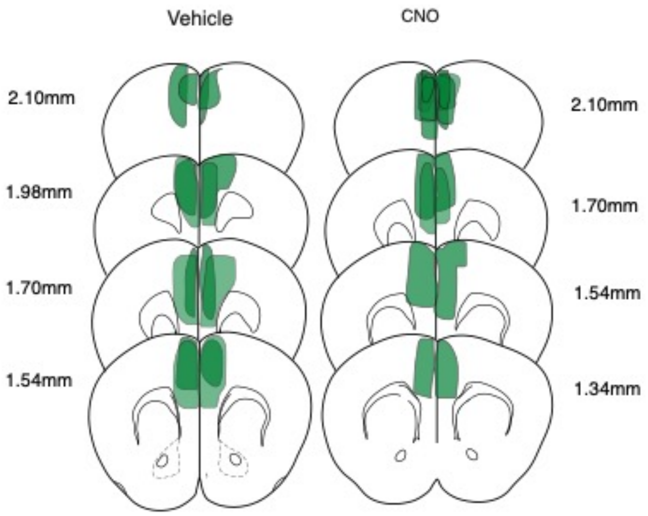
Vehicle
CNO
2.10mm
1.98mm
1.70mm
1.54mm
2.10mm
1.70mm
1.54mm
1.34mm

B

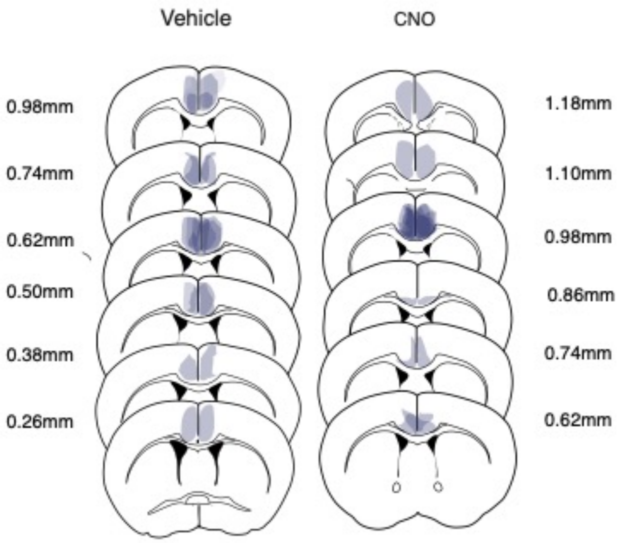
Vehicle
CNO
0.98mm
0.74mm
0.62mm
0.50mm
0.38mm
0.26mm
1.18mm
1.10mm
0.98mm
0.86mm
0.74mm
0.62mm

**Supplementary Figure 2. AAV-Syn::mCherry-2A-HA-hM4Di virus diffusion for the recent and remote consolidation experiments.** Schematic representation of mCherry protein expression in vmPFC **(A),** and in the aCC **(B)**. mCherry maximum extension is represented in green for each single mouse for the vmPFC group and in blue for each single mouse of aCC group. Coordinates are expressed as mm mm anterior to bregma bregma.

A

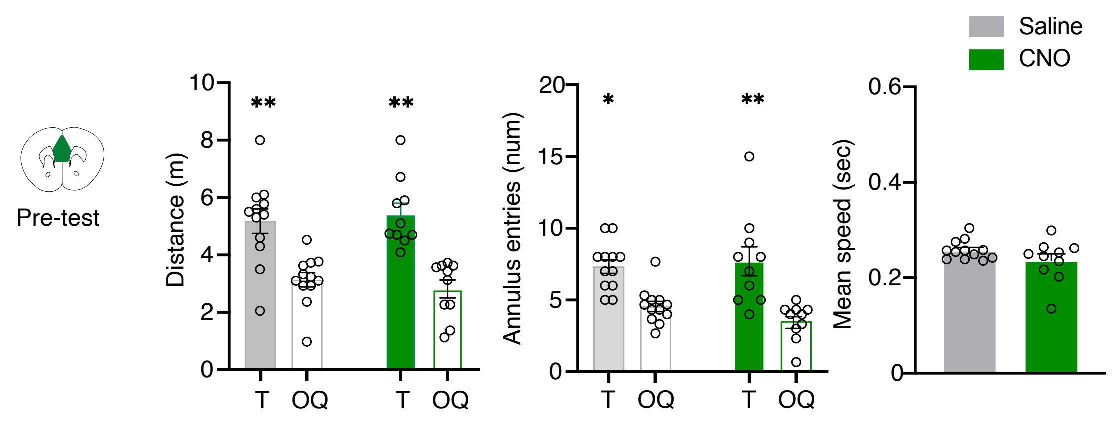

Pre-test
Distance (m)
T
OQ
Annulus entries (num)
Mean speed (sec)
Saline
CNO


B

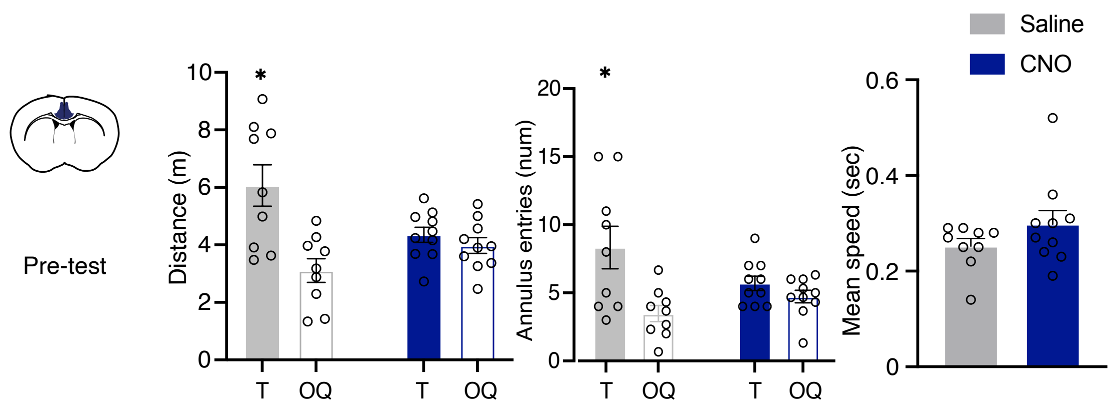

Pre-test
Distance (m)
T
OQ
Annulus entries (num)
Mean speed (sec)
Saline
CNO

**Supplementary Figure 3. Effects of pre-test loss-of-function manipulations of the vmPFC or the aCC on probe test at remote time points. (A) (left panel)** Mean distance travelled (m) ± SEM on probe test for vmPFC group [ANOVA of quadrant $F_{(1,20)}$= 43.67, p<0.0001; treatment $F_{(1,20)}$= 0.016, p= 0.919; session x treatment $F_{(1,20)}$= 0.6562, p= 0.427]. **(center panel)** Mean annulus entries ± SEM on probe test for vmPFC group [ANOVA of quadrant $F_{(1,20)}$= 27.76, p<0.0001; treatment $F_{(1,20)}$= 0.394, p= 0.537; session x treatment $F_{(1,20)}$= 1.11, p= 0.304]. **(right panel)** Mean of speed (sec) on probe test for vmPFC group [$t_{(20)}$= 1.503, p= 0.14]. **(B) (left panel)** Mean distance travelled (m) ± SEM on probe test for aCC group [ANOVA of quadrant $F_{(1,17)}$= 10.54, p<0.0004; treatment $F_{(1,17)}$= 0.0159, p= 0.901; session x treatment $F_{(1,20)}$= 2.872, p= 0.108]. **(center panel)** Mean annulus entries ± SEM on probe test for aCC group [ANOVA of quadrant $F_{(1,17)}$= 9.61, p<0.0006; treatment $F_{(1,17)}$= 0.381, p= 0.544; session x treatment $F_{(1,20)}$= 4.126, p= 0.058]. **(right panel)** Mean of speed (sec) on probe test for aCC group [$t_{(17)}$= 1.343, p= 0.19]. Target quadrant (T); Other Quadrants (OQ). ** p≤ 0.01; * p≤ 0.05 vs target quadrant (within group).

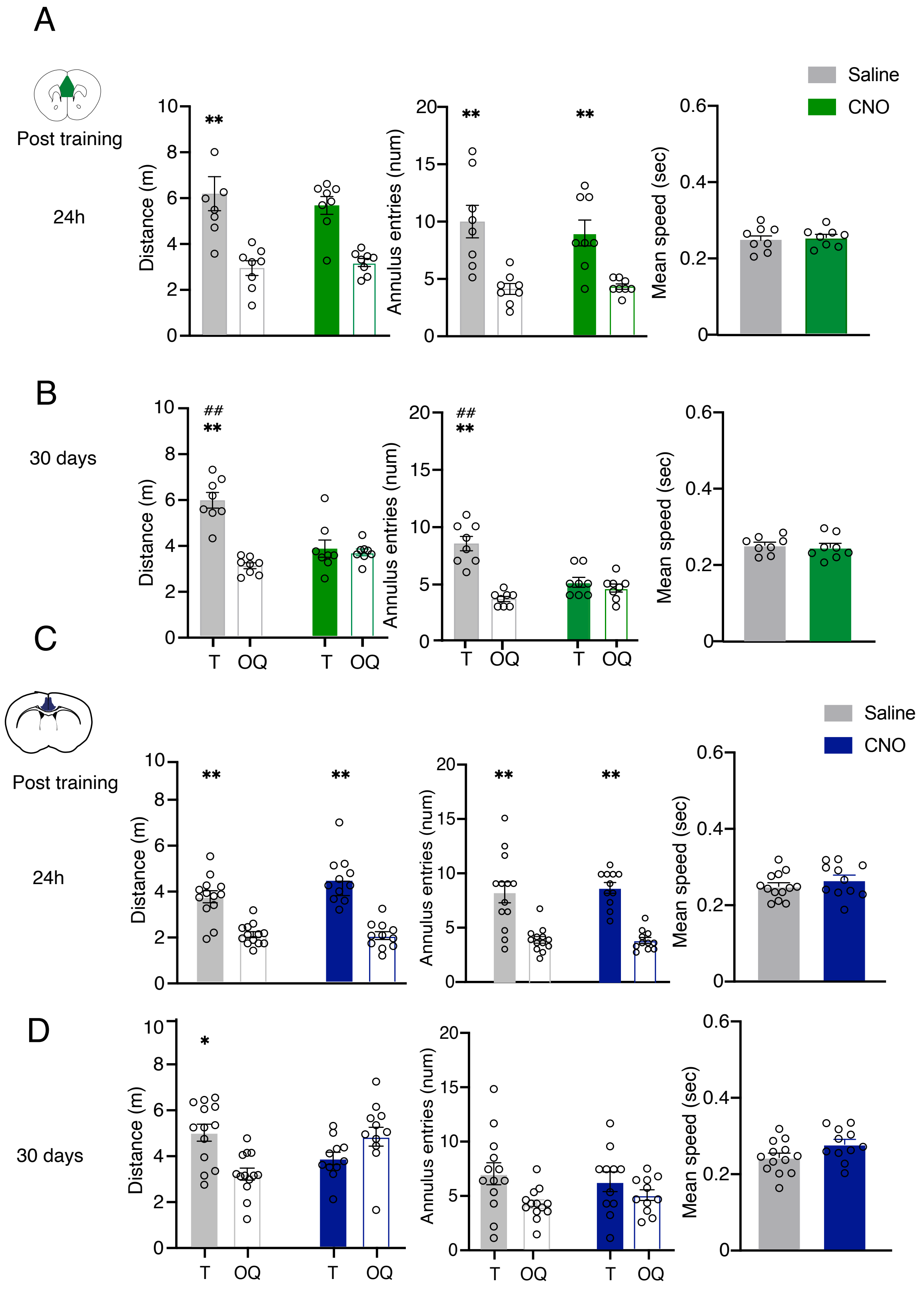
A
Post training
24h
Saline
CNO
Distance (m)
Annulus entries (num)
Mean speed (sec)
**
B
30 days
##
T
OQ
C
Post training
24h
D
30 days
*

**Supplementary Figure 4. Effects of immediate post-training loss-of-function manipulations of the vmPFC or the aCC on probe test at recent (24h) and remote (30d) time points. (A)** Probe trial 24 hour after training of mice infected in the vmPFC and administered immediately post-training with vehicle or CNO. **(left panel)** Mean of distance travelled (m) ± SEM [ANOVA of quadrant $F_{(1,14)}$= 26.82, p<0.0001; treatment $F_{(1,14)}$= 0.158, p= 0.696; session x treatment $F_{(1,14)}$= 0.46, p= 0.506]; (**center panel**) Mean annulus entries ± SEM [ANOVA of quadrant $F_{(1,14)}$= 22.84, p<0.001; treatment $F_{(1,14)}$= 0.277, p= 0.606; session x treatment $F_{(1,14)}$= 0.300, p= 0.592]; (**right panel**) Mean of speed (sec) ± SEM [$t_{(14)}$= 1.979, p= 0.84] on probe test 24h after training for vmPFC infected groups. **(B).** Probe test 30 days after training of vmPFC infected mice administered with vehicle or CNO immediately after training. (**left panel**) Mean of distance travelled (m) ± SEM [ANOVA of quadrant $F_{(1,14)}$= 34.38, p=0.00017; treatment $F_{(1,14)}$= 7.16, p= 0.018; session x treatment $F_{(1,14)}$= 27.42, p= 0.00013]; (**center panel**) Mean annulus entries ± SEM [ANOVA of quadrant $F_{(1,14)}$= 36.38, p<0.001; treatment $F_{(1,14)}$= 7.92, p=0.014; session x treatment $F_{(1,14)}$= 24.02, p<0.001]. (**C**) Effect of immediate post-training administrations of vehicle or CNO on probe trial 24 hours after training in aCC infected mice**. (left panel**) Mean of distance travelled (m) ± SEM [ANOVA of quadrant $F_{(1,22)}$= 63.75, p<0.001; treatment $F_{(1,22)}$= 3.21, p=0.087; session x treatment $F_{(1,22)}$= 2.405, p=0.135]; (**center panel**) Mean annulus entries ± SEM [ANOVA of quadrant $F_{(1,22)}$= 52.07, p<0.001; treatment $F_{(1,22)}$= 0.09, p=0.765; session x treatment $F_{(1,22)}$= 0.127, p=0.724]; (**D)** Probe trial 30 days after training of aCC infected mice administered immediately post-training administrations with vehicle or CNO. **(left panel**) Mean of distance travelled (m) ± SEM [ANOVA of quadrant $F_{(1,22)}$= 13.76, p=0.0012; treatment $F_{(1,22)}$= 1.55, p=0.226; session x treatment $F_{(1,22)}$= 0.78, p=0.38]; (**center panel**) Mean annulus entries ± SEM [ANOVA of quadrant $F_{(1,22)}$= 5.16, p=0.033; treatment $F_{(1,22)}$= 0.0002, p=0.99; session x treatment $F_{(1,22)}$= 0.71, p=0.408]; (**right panel**) Mean of speed (sec) ± SEM [$t_{(22)}$= 1.993, p= 0.058]. Target quadrant (T); Other Quadrants (OQ). *p≤ 0.05; ** p≤ 0.01 vs target quadrant (within group); ## p≤ 0.01 target vs target quadrant (between groups) (Tukey, HSD).
